# Structural Dynamics of the Ubiquitin Specific Protease USP30 in Complex with a Cyanopyrrolidine-Containing Covalent Inhibitor

**DOI:** 10.1101/2024.07.20.604388

**Authors:** Darragh P O’Brien, Hannah BL Jones, Yuqi Shi, Franziska Guenther, Iolanda Vendrell, Rosa Viner, Paul E Brennan, Emma Mead, Tryfon Zarganes-Tzitzikas, John B Davis, Adán Pinto-Fernández, Katherine S England, Emma J Murphy, Andrew P Turnbull, Benedikt M Kessler

## Abstract

Inhibition of the mitochondrial deubiquitinating enzyme USP30 is neuroprotective and presents therapeutic opportunities for the treatment of idiopathic Parkinson’s Disease and mitophagy-related disorders. We have integrated structural and quantitative proteomics with biochemical assays to decipher the mode of action of covalent USP30 inhibition by a small molecule containing a cyanopyrrolidine reactive group, **USP30-I-1**. The inhibitor demonstrated high potency and selectivity for endogenous USP30 in neuroblastoma cells. Enzyme kinetics and Hydrogen Deuterium eXchange mass spectrometry (HDX-MS) infers that the inhibitor binds tightly to regions surrounding the USP30 catalytic cysteine and positions itself to form a binding pocket along the thumb and palm domains of the protein, thereby interfering its interaction with ubiquitin substrates. A comparison to a non-covalent USP30 inhibitor containing a benzosulfonamide scaffold revealed a slightly different binding mode closer to the active site Cys77, which may provide the molecular basis for improved selectivity towards USP30 against other members of the DUB enzyme family. Our results highlight advantages in developing covalent inhibitors, such as **USP30-I-1**, for targeting USP30 as treatment of disorders with impaired mitophagy.

## INTRODUCTION

Ineffective repair or clearance of damaged mitochondria negatively affects cellular health and is an integral feature of several neurodegenerative disorders such as Parkinson’s Disease (PD), Alzheimer’s Disease, and amyotrophic lateral sclerosis, in addition to cardiomyopathy and ageing ^1–3^. Without their elimination, defective mitochondria accumulate, resulting in the excessive production of highly reactive and toxic oxygen species ^4, 5^. This results in extensive damage to overall cellular integrity and survival. Mitophagy, the cell’s specialized quality control system for the clearance of damaged mitochondria, can proceed through the concerted action of two enzymes; the mitochondrial outer membrane (MOM)-associated ubiquitin (Ub) serine/threonine kinase PINK1, and the cytoplasmic E3 ligase Parkin ^6^. PINK1 and Parkin activity leads to the hyper-ubiquitination of damaged proteins on the MOM, providing the molecular signal for their removal. Mitophagy is negatively regulated by several deubiquitinating (DUB) enzymes, including Ubiquitin Specific Proteases (USP) 8, 14, 15, 30, and 35 ^7–11^. The central action of these DUB enzymes is to remove Ub moieties from mitochondrial proteins and, as a consequence, modulate mitophagy. Of these, USP30 has been associated with mitophagy, likely due to its exclusive expression on the MOM ^12^. Aside from mitochondria, its analogous expression on peroxisomes widely implicates it in pexophagy and redox homeostasis ^13, 14^.

An accumulation of damaged mitochondria has been linked to both familial and sporadic forms of PD. Loss-of-function mutations in PINK1 and PRKN genes lead to a build-up of defective mitochondria and a gradual loss of dopaminergic neurons in the basal ganglia, resulting in a rare and inherited form of early-onset Parkinsonism ^15, 16^. Inhibiting USP30 may therefore boost mitochondrial turnover in both early onset Parkinsonism and in PD, at least in this population of patients, offering a new strategy for the treatment of the disease and similar neurodegenerative conditions. This has justifiably attracted a great deal of attention in both academia and industry, with several small-molecule inhibitors, which can chemically reduce USP30 activity currently in development ^17–22^. Importantly, USP30 inhibitors MTX652 and MTX325 are being tested in clinical trials for the treatment of acute kidney injury (AKI) and PD ^23^.

We have recently described the structural interplay of USP30 following complex formation with a non-covalent benzosulfonamide inhibitor “**USP30_inh_**”, which has been shown to boost mitophagy in dopaminergic neurons by reducing USP30 activity ^19, 22, 24^. A combination of Activity-Based Protein Profiling Mass Spectrometry (ABPP-MS), Bio-layer Interferometry, Hydrogen/Deuterium eXchange Mass Spectrometry (HDX-MS) and computational modeling allowed us to comprehensively profile the potency, selectivity, and mechanism of inhibition of **USP30_inh_** for dampening endogenous USP30. This was the first study of its kind to provide detailed structural and mechanistic information on the interaction of USP30 with any active small-molecule drug targeting it.

We now extend our study to structurally profile USP30’s interaction with a small *covalent* cyanopyrrolidine scaffold-containing inhibitor, **USP30-I-1** (patent WO2020212350A1; Mission Therapeutics) ^25, 26^. Covalent inhibitors possess significant advantages over their non-covalent counterparts, including a more prolonged duration of action and the formation of more specific and irreversible bonds with their substrates, enhancing overall drug efficacy ^27^. Newer-generation USP30 inhibitors are therefore likely to harness covalent bond formation between compound and protein. Using a biophysical and structural proteomics approach similar to our previous work ^24^, we show that **USP30-I-1** is highly selective and potent against endogenous USP30 when compared to the >40 other endogenous DUBs identified in the neuroblastoma-derived SH-SY5Y cell line. Covalent **USP30-I-1** binds to USP30 with a similar affinity to the previously characterized non-covalent **USP30_inh_**, with binding primarily restricted to a small region covering the USP30 active site cysteine residue. Several regions along the palm and thumb domains of the protein also become solvent protected in the presence of **USP30-I-1**. Directly comparing the two studies, we decipher commonalities and differences between covalent and non-covalent mechanisms of USP30 inhibition. Our biochemical, structural, computational, and biophysical data ultimately empowers the design of next-generation USP30 inhibitors to further drive drug discovery campaigns in the USP30 inhibitor space.

## EXPERIMENTAL SECTION

### USP30 INHIBITOR PURITY

**USP30-I-1** has a cyanopyrrolidine warhead for covalent reaction with Cys77 of USP30. The compound was 93% pure at 254 nm UV and 100% by ELSD (**Figure S1**). USP30 inhibitor **USP30-I-1** synthesis and characterization was reported previously (patent WO2020212350A1; Mission Therapeutics) ^23, 25, 26^.

## 1. ABPP ASSAY

### Cell culture and lysis

SH-SY5Y cells were cultured and lysed as previously reported ^24^. Briefly, cells were cultured at 37°C, 5% CO_2_ in Eagle’s minimum essential medium and Ham’s F12 nutrient mix supplemented with fetal bovine serum (FBS), non-essential amino acids, and Glutamax. Cells were washed with phosphate buffered saline (PBS), scraped, and collected by centrifugation at 200 x g. Pellets were resuspended in 50 mM Tris base, 5 mM MgCl_2_.6 H_2_O, 0.5 mM EDTA, 250 mM sucrose, 1 mM DTT (pH 7.5) and lysed with glass bead beating. After 4°C clarification at 600 x g for 10 min, lysate protein concentrations were measured by bicinchoninic acid assay.

### HA-Ub-PA activity-based probe profiling

HA-Ub-PA was synthesized as outlined previously ^28, 29^. Methodology for HA-Ub-PA activity-based probe profiling was described in our previous study on a USP30 non-covalent inhibitor ^24^. Briefly, SH-SY5Y lysates were incubated for 1 hr at 37°C with either **USP30-I-1** or dimethyl sulfoxide (DMSO) at the indicated concentrations in duplicate. Labeling with HA-Ub-PA was then carried out at 37°C for 45 min. The reaction was then quenched with SDS and NP-40 and diluted in 50 mM Tris, 0.5% NP-40, 150 mM NaCl, and 20 mM MgCl_2_.6 H_2_O, pH 7.4. HA-Ub-PA-bound DUBs were immunoprecipitated overnight at 4°C with end-over-end rotation using 150 µL anti-HA agarose slurry. HA-Ub-PA DUB complexes were eluted with 2x Laemmli buffer, reduced, alkylated and cleaned up for LC-MS/MS using S-Trap™ micro columns ^30^. Samples were digested with 2 µg of trypsin. Eluates were dried and resuspended in 0.1 % formic acid (FA).

### Western blot quantitation

Densitometry image analysis was applied for western blot quantification of the unbound and HA-Ub-PA bound USP30 bands. The percentage of signal from the HA-Ub-PA bound USP30 band relative to signal from both bands was used to quantify USP30 activity and inhibition. Activity was normalized relative to the negative (no HA-Ub-PA) and positive (HA-Ub-PA) controls. The IC_50_ was extracted from fitting **USP30-I-1** inhibition quantitation with the equation: Y=100/(1+10^(X-LogIC_50_)) in Prism (version 10.1.1).

### LC-MS/MS data collection

Samples were analyzed by LC-MS/MS using the Vanquish Neo UHPLC (Thermo) connected to Thermo Orbitrap Ascend mass spectrometer (Thermo). The Vanquish Neo was operated in “Trap and Elute” mode using a PepMap™ Neo trap (185 μm, 300 μm x 5 mm) and an EASY-Spray™ PepMap™ Neo column (50 cm x 75 μm, 1500 bar). Tryptic peptides were separated over a 60 min linear gradient of 3 to 20% B – 80% ACN, 0.1% FA in 40 min and 20 to 35% B in 20 min. The system was maintained at a 300 nL/min flow rate. Samples were analyzed by tandem mass spectrometry as previously described ^24, 31, 32^. Data were collected on an Orbitrap Ascend Tribrid mass spectrometer (Thermo Scientific) over *m/z* 350–1650 by Data Independent Acquisition (DIA), with a 50K resolution, a maximum injection time of 91 msec, an AGC set to 125%, and a radio frequency lens set to 30%. MS2 data were collected using the tMSn scan function, with 40 DIA scan windows of variable widths, an Orbitrap resolution of 30K, a normalized AGC target of 1000%, a maximum injection time set to auto, and a collision energy set to 30%.

### LC-MS/MS data processing

LC-MS/MS data was processed using DIA-NN (version 1.8.1) for a library free search using a *Homo sapiens* Uniprot database (20,416 entries, retrieved February 15^th^, 2023) with all settings left as default ^33^. The unique gene matrix output was used to identify DUBs that were HA-Ub-PA enriched, with intensities 5-fold higher in the HA-Ub-PA positive control when compared to the negative control with no HA-Ub-PA. YOD1 and USP9Y were removed from analysis as they were not consistently identified across replicates. Significant inhibition of DUBs was identified using two-way ANOVA analysis in Prism (version 10.1.1) with the Dunnett method for multiple correction testing (\*\*\*\**p* < 0.0001).

## 2. ENZYME KINETICS

### USP30 Activity Assay

Fluorescence intensity measurements were used to monitor the cleavage of a ubiquitin-rhodamine substrate. All activity assays were performed in black 384-well plates in assay buffer (20 mM Tris-HCl, pH 8.0, 150 mM Potassium Glutamate, 0.1 mM TCEP and 0.03% Bovine Gamma Globulin) with a final assay volume of 20 μL. A concentration of 0.2 nM USP30 (residues 64-502Δ179-216 & 288-305, Viva Biotech (Shanghai) Ltd.) was added and preincubated with **USP30-I-1** for 30 min. A total of 25 nM ubiquitin-rhodamine 110 (Ubiquigent) was added to initiate the reaction and the fluorescence intensity was recorded for 30 min on a PherastarFSX (BMG Labtech) with an Ex485/Em520 optic module. Initial rates were plotted against compound concentration to determine the IC_50_.

### Kinetic assays – determination of k_inact_/K_i_

Kinetic assays were performed in a 384-well Sensoplate™ in assay buffer with a final assay volume of 50 μL. The assay was carried out using 5 nM USP30 and reactions were started by simultaneous addition of 180 nM ubiquitin-rhodamine 110 to all 384 wells using the FLIPR® Tetra (Molecular Devices) dispense function. Fluorescence was monitored every 3 sec (excitation wavelength 470-495 nm, emission 515-575 nm, camera gain 70, exposure time 0.6 sec, excitation intensity 80%) over 10 min.

Analysis was performed in GraphPad Prism. Fluorescence intensity was plotted vs. inhibitor concentration at each time point to determine IC_50_, IC_50_ was then plotted vs. incubation time and fitted to Eq. 1 (Krippendorf) ^34^ to obtain the inhibition constant K_I_ and the rate of enzyme inactivation k_inact,_ the values can be used to obtain k_inact_/K_I_, a second order rate constant describing the efficiency of covalent bond formation.

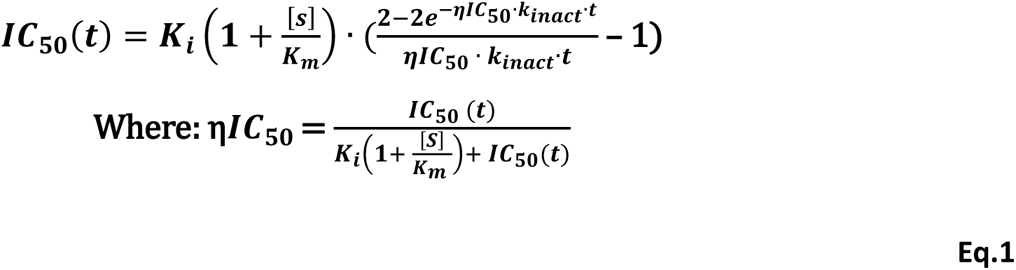

K_inact_/K_I_ was also determined using the traditional method of fitting the progress curves to Eq. 2 and plotting k_obs_ vs [**USP30-I-1**] and fitting with Eq. 3 (supplemental).

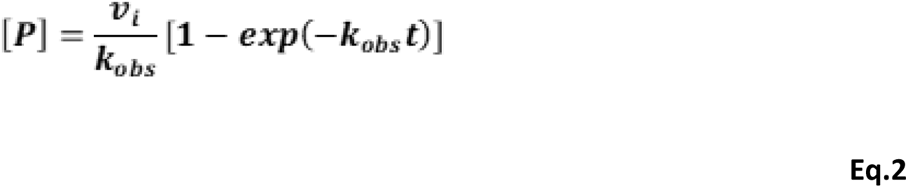

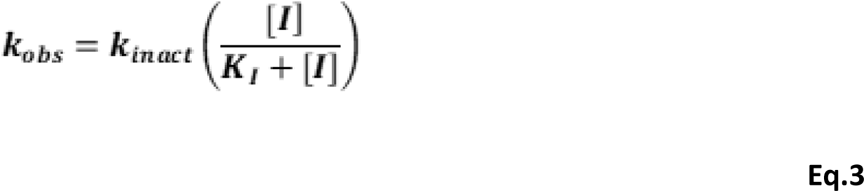

## 3. HDX-MS

### Materials

The same recombinant USP30 construct (residues 64-502Δ179-216 & 288-305, Viva Biotech (Shanghai) Ltd.) that was used in our *in vitro* enzyme kinetics analysis was also used for HDX-MS. LC-MS grade water, LC-MS grade 0.1% FA, LC-MS grade ACN with 0.1% FA in water were purchased from Fisher Scientific (Hampton, NH). Guanidine hydrochloride 8M solution, TCEP-HCl, and DMSO were purchased from Thermo Scientific (Rockford, IL). Citric acid, Sodium Chloride and HEPES were purchased from Sigma-Aldrich (St. Louis, MO). Deuterium oxide (99+ %D) was purchased from Cambridge Isotope Laboratories (Tewksbury, MA).

### Sample preparation

Initial stock concentrations of USP30 and compound **USP30-I-1** in DMSO were 66 µM and 10 mM, respectively. For in-solution HDX-MS, a working sample of USP30 and **USP30-I-1** (*i.e*., the holo-USP30) was prepared by volume-to-volume mixture at a molar ratio of 1:2 (USP30:**USP30-I-1)** and diluted to a nominal concentration of 11 μM for the USP30, and 22 μM for **USP30-I-1**. The reference state (*i.e*., apo-USP30) was 11 μM USP30 protein which was supplemented with DMSO in place of the compound.

### HDX LC-MS/MS data collection

In-solution HDX-MS was performed as follows: the labeling buffer was 50 mM HEPES pD 7.2, 400 mM NaCl, 2 mM TCEP in D_2_O, and the quench buffer was 2M Guanidine HCl, 100 mM citric acid, pH 2.3 in water. The pH of the labeling buffer was measured and corrected to pD (pD=pH+0.4). Approximately, 3.5 µL of the USP30 with **USP30-I-1** complex mixture was diluted with the labeling buffer (1:20 ratio) and incubated in D_2_O buffer at 20ᵒC for 30, 60, 600, and 3600 sec in triplicates. Non-deuterated controls were prepared in an identical manner, except H_2_O was used in place of D_2_O in the labeling step. Then, the labeled sample was quenched by adding quench buffer (1:1 ratio) and incubated for 180 sec. LC/MS bottom-up HDX was performed using a Thermo Scientific™ Ultimate™ 3000 UHPLC system and Thermo Scientific^™^ Orbitrap Exploris^™^ 480 Hybrid^™^ mass spectrometer. The quenched samples (50 µL) were digested with a pepsin/proteaseXIII (NovaBioAssays, MA) column (2.1 x 3.0 mm) at 8ᵒC for 2 min and then trapped in a 1.0 mm x 5.0 mm, 5.0 µm trap cartridge (Thermo Scientific™ Acclaim PepMap100) for desalting. Peptides were separated on a Thermo Scientific™ Hypersil Gold^™^, 50 x 1 mm, 1.9 um, C18 column with a linear gradient of 10% to 40% Buffer B (A: water, 0.1% FA; B: ACN, 0.1% FA) at a flow rate of 40 µL/min. Pepsin wash was performed for each run to limit carry-over. To limit back-exchange, the quenching, trapping, and separation steps were performed at near 0ᵒC. Labeling, quenching, and online digestion were performed in a fully automated manner with Chronnect HDX workstation by Trajan.

### Data analysis

Before conducting HDX-MS experiment, an unspecific digested peptide database was created for non-deuterated USP30 sample using data-dependent and targeted HCD-MS^2^ acquisition. Peptide identification was performed using BioPharma Finder (v5.1). HDX-MS data files were processed and manually curated with the USP30 peptide database using HDExaminer by Trajan. A single charge state with high quality spectra for all replicates across all HDX-MS labeling times was chosen to represent HDX for each peptide. A hybrid statistical significance approach was performed afterwards using an in-house MATLAB script^35^. The significant differences observed at each residue was used to map HDX-MS consensus effects (based on overlapping peptides) onto the catalytic domain using the X-ray structure of human USP30 in complex with a Fab fragment antibody and covalent inhibitor, **552**, as the template (PDB code: 8D1T; unpublished).

## 4. MOLECULAR DOCKING

We performed molecular docking simulations using the covalent docking method in ICM-Pro (MolSoft LLC). There are two inhibitor-bound X-ray structures for human USP30 in the Protein Data Bank, corresponding to the catalytic domain in complex with a Fab fragment antibody and covalent inhibitors **552** (PDB code: 8D1T; unpublished) and **829** (PDB code: 8D0A; unpublished), at 2.94 Å and 3.19 Å resolution, respectively. The highest resolution human USP30 catalytic domain inhibitor complex structure (corresponding to PDB code: 8D1T) in which the Fab fragment antibody and covalent inhibitor, **552**, had been removed, was used as the target receptor for docking studies with **USP30-I-1**. The *in-silico* binding pose of **USP30-I-1** with the best docking score is shown in **Figure 3**.

## RESULTS AND DISCUSSION

### USP30-I-1 targets endogenous USP30 in a highly potent and selective manner

We performed ABPP-MS in SH-SY5Y neuroblastoma cell lysates to determine the potency and selectivity of **USP30-I-1,** a substituted cyanopyrrolidine derivative (**Figure 1A**). Endogenous USP30 inhibition was confirmed via the prevention of HA–Ub–PA-USP30 binding by **USP30-I-1** in a concentration-dependent fashion (**Figure 1B**). **USP30-I-1** was observed to be a potent inhibitor of USP30, with an IC_50_ of 94 nM (**Figures 1C**). Our quantitative MS analysis showcased the high selectivity of **USP30-I-1** for USP30, with no significant activity against any of the other 40 endogenous DUBs detected in SH-SY5Y extracts at the lowest inhibitor concentrations (**Figures 1D** and **1E**). At higher concentrations of 10 µM, however, we observed that **USP30-I-1** reduces the HA-Ub-PA labeling of a number of other DUBs (**Figure S2**). This included approximately 50% inhibition of USP10. However, as USP10 is not strongly labelled by HA-Ub-PA, the inhibitory profile of USP10 by **USP30-I-1** could not be validated by western blot (**Figure S3A**). Moreover, **USP30-I-1** is far more potent for USP30 than it is for USP10 (**Figure S3B**). In conclusion, ABPP-MS demonstrated that **USP30-I-1** is a selective and potent inhibitor of USP30 at concentrations ≤1 µM.

**Figure 1.**
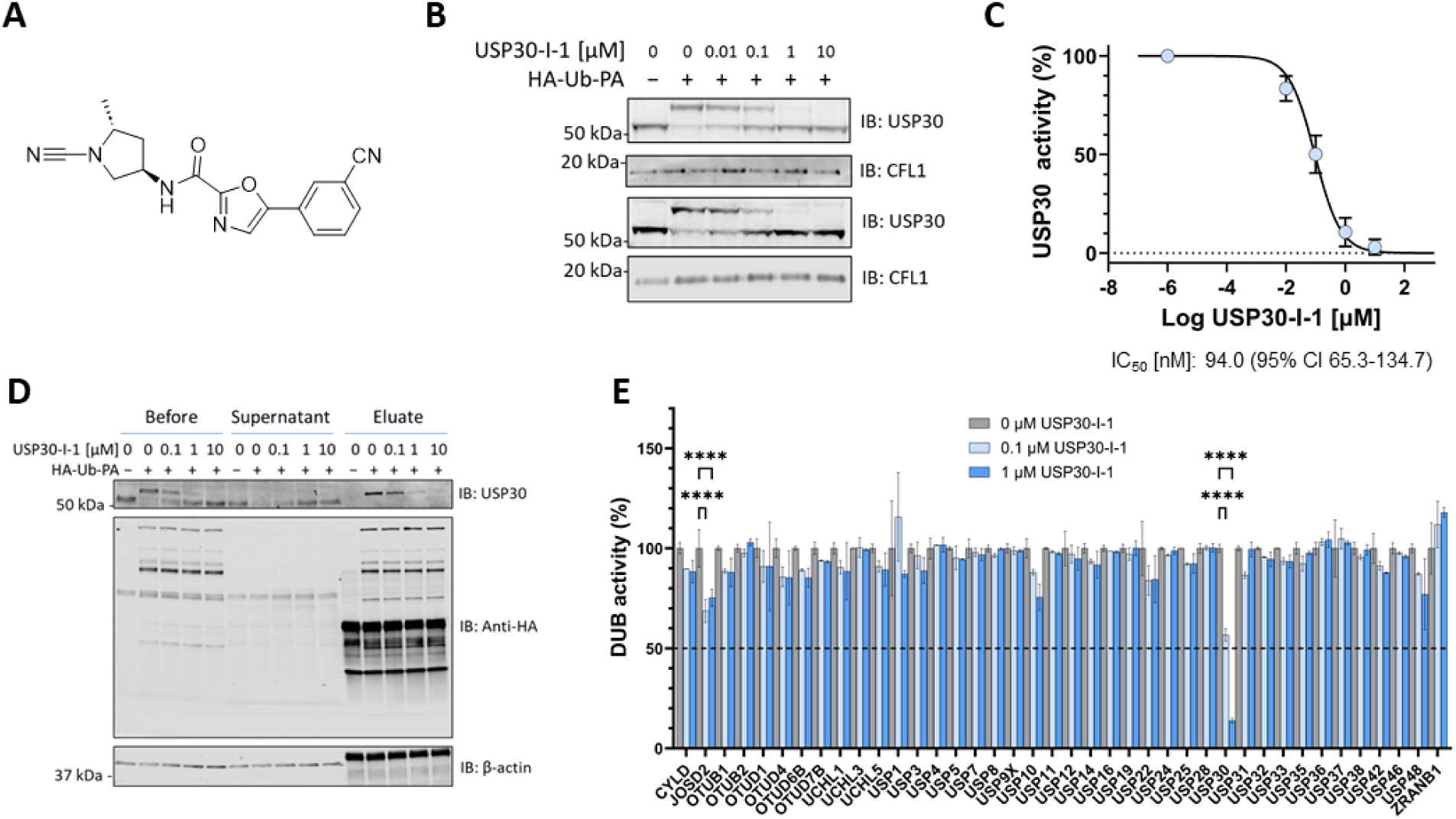
USP30-I-1 is selective and potent for endogenous USP30. (**A**) Structure of **USP30-I-1** (**B, C**) Immunoprecipitation of HA-Ub-PA DUB complexes, with USP30 HA-Ub-PA binding and immunoprecipitation reduced by **USP30-I-1** in a concentration dependent manner. (**D**) Anti-HA blot showing no inhibition of HA-Ub-PA binding for other DUBs. (**E**) LC-MS/MS quantitation of HA-Ub-PA enriched DUBs with **USP30-I-1** at indicated concentrations relative to positive control (****p < 0.0001).

We then undertook *in vitro* biochemical assays to determine the full enzyme kinetics of **USP30-I-1**. In the first instance, fluorescence intensity measurements were used to monitor the interaction of a fluorogenic Ub-rhodamine substrate with a recombinant, truncated version of USP30 ^36^, with and without inhibitor. **USP30-I-1** inhibited USP30 in a dose-dependent manner, with an IC_50_ of ∼4 nΜ when USP30 was preincubated with inhibitor for 30 min. (**Figure 2A**). The lower IC_50_ value observed for **USP30-I-1** in the *in vitro* work was likely a result of reduced non-specific inhibitor occlusion as compared to the cellular matrix, and a similar phenomenon was observed in our **USP30**_inh_ work ^24^. Rates of inhibition determined from the Ub–rhodamine cleavage progress curves (**Figure 2B**) without USP30 preincubation, were plotted against [**USP30-I-1**] (**Figure 2B** Inset) to determine k_inact_/K_I_ (traditional method). IC_50_ values were also measured and plotted against incubation time to calculate k_inact_/K_I_ (Krippendorf method) (**Figures 2D** and **2E**), with both forms of analysis giving comparable parameters (**Figure 2E**). To conclude, **USP30-I-1** binds to USP30 in a tight manner and displays kinetic properties consistent with covalent attachment.

**Figure 2.**
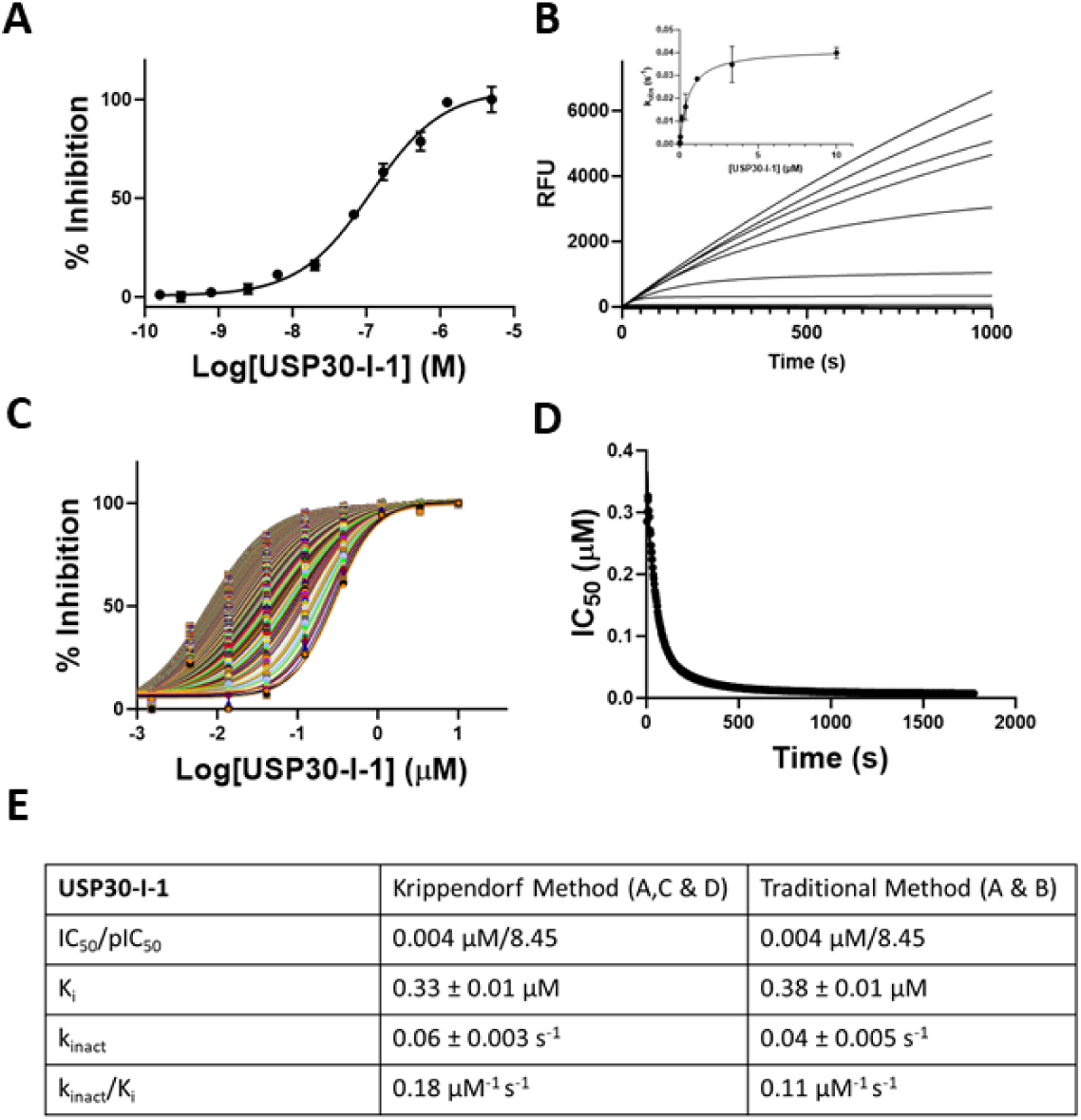
Covalent USP30-I-1 tightly binds to recombinant USP30. (**A**) Dose dependent inhibition of USP30 by **USP30-I-1**. (**B**) Progress curves recorded on the FLIPR**^®^** Tetra. Traditional method for determining kinetic constants associated with covalent binders. k_obs,_ determined by fitting the progress curves to Eq. 2 (Methods), is plotted vs. [Compound] and fitted to Eq. 3. (Methods) to determine K_I_ and k_inact_. (**C**) Krippendorf method used for determining covalent kinetic constants. Time-dependent IC_50_ curves. Each curve represents inhibition data at an individual incubation time from 3-600 s. (**D**) IC_50_ values vs. incubation time fitted to Eq. 1. (Methods) to obtain K_I_ and k_inact_. (**E**) Data table of kinetic constants.

### USP30-I-1 binds to the catalytic cysteine of USP30, inducing conformational changes in the active site

We used HDX-MS to determine the conformational dynamics associated with the interaction between **USP30-I-1** and USP30, in addition to pinpointing its precise location of small-molecule binding. From in excess of 700 unique peptide identifications, 161 peptides were used in the final data analysis, due to the ability to confidently detect each of them across all HDX-MS labeling time points. This resulted in an overall sequence coverage of 98% for the recombinant USP30 construct, which had an average redundancy of 4.71 peptides covering each amino acid. When directly comparing apo-USP30 to holo-USP30, the majority of the USP30 protein sequence had no significant differential deuterium uptake in the presence of **USP30-I-1**, confirming that, as one may expect upon interaction of a small-molecule to a much larger protein, binding was restricted to only a small portion of USP30 itself (**Figure 3A**). The hybrid statistical analysis comparing the apo– and holo-states easily isolated a handful of peptides/labeling time points that became significantly shielded from the deuterium buffer in the presence of the compound, indicative of binding and/or interaction. These primarily matched to peptides covering the sequence ^70^LVNLGNTCF^78^, which comprises the catalytic Cys77 and preceding loop, and where a mean perturbation in deuterium labeling between states of up to 60% was observed across all time points. It should be noted, however, that solely in the holo-state, the overall peptide quality was reduced in this region of USP30, presumed to a direct result of the strong covalent attachment of the inhibitor to the peptide under HDX-MS experimental conditions. This peptide contains the catalytic Cys77 covalently modified with the inhibitor and may therefore have different ionization properties. Nevertheless, this observation provides an additional layer of confidence that the catalytic Cys77 is indeed the primary binding site of **USP30-I-1** on the protein. We subsequently mapped all HDX-MS behaviours onto the structure of human USP30 catalytic domain using the X-ray structure of human USP30 catalytic domain in complex with the covalent inhibitor, **552**, and a Fab fragment antibody, in which the covalent inhibitor and Fab fragment antibody had been removed, as the template (**Figure 3B**). This structure is representative of the catalytic domain conformation in complex with a covalent inhibitor as exemplified by both covalent inhibitor structures present in the Protein Data Bank. Our mapping further highlighted the strong interaction of **USP30-I-1** with the region encompassing the catalytic Cys77 (highlighted in magenta in **Figure 3B**), but also supported the identification of other regions of human USP30 catalytic domain (highlighted in red) that become solvent protected, albeit to a much lesser extent. These include the sequences ^150^YRWQISSF^157^ (corresponding to the switching loop), ^322^CIHLQRLSWSSHGTPLKRH^340^ (corresponding to blocking loop 1), ^446^GDMHSGHFVTY^456^ (corresponding to blocking loop 2), and ^465^NPLSTSNQWL^474^ (residues preceding and forming part of the β-strand that accommodates catalytic residue, Ser477, in the central β-sheet in the palm subdomain) (**Figures 3B** and **3C**). In conclusion, our differential HDX-MS allowed us to pinpoint regions of USP30 which are involved in **USP30-I-1** binding.

**Figure 3.**
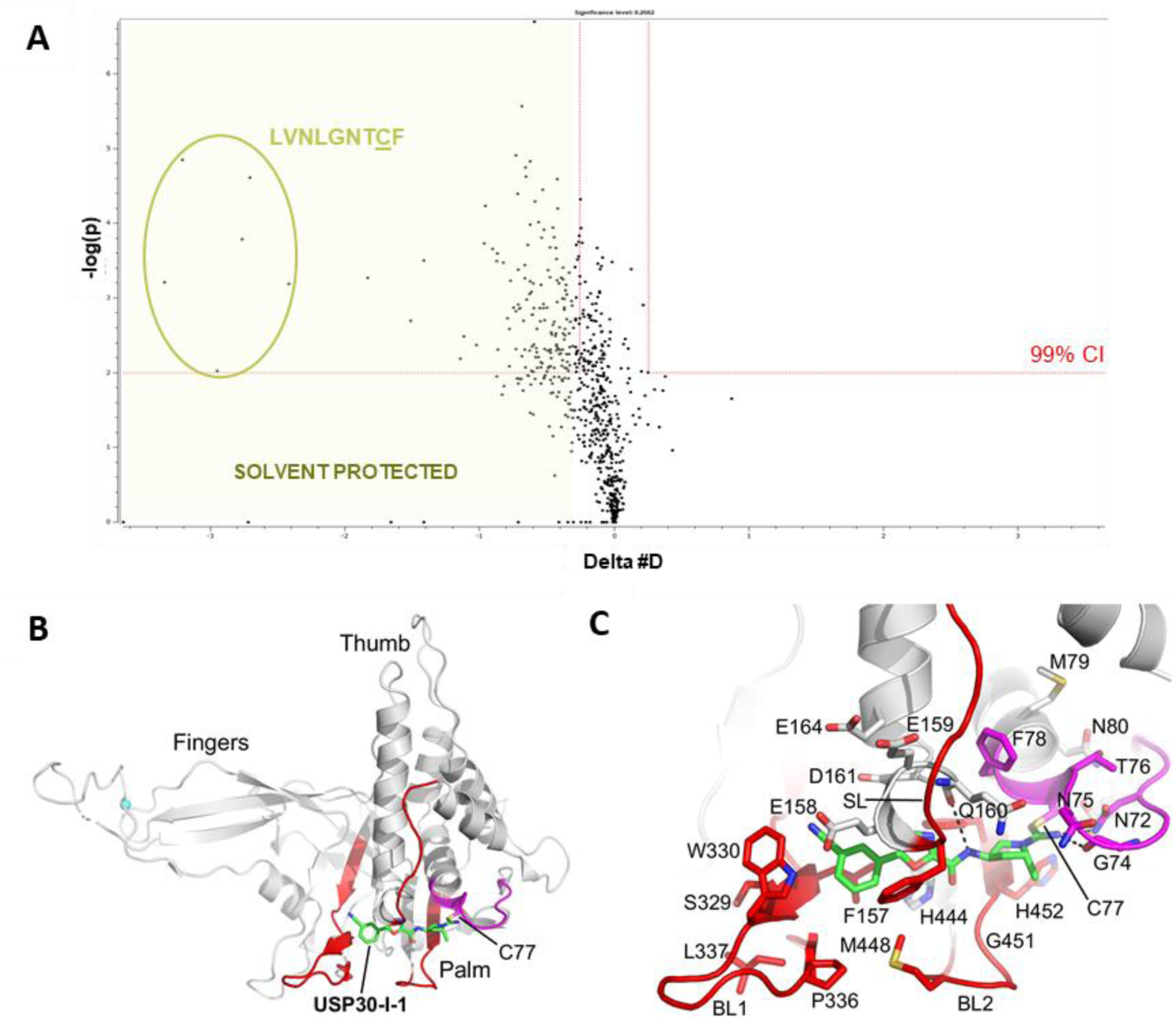
USP30-I-1 binds to the catalytic cysteine of USP30. (**A**) HDX-MS shows that the USP30 region ^70^LVNLGNNTCF^78^ which surrounds the catalytic Cys77 (underlined) is significantly solvent protected in the presence of **USP30-I-1.** This implies that it is the primary location of compound binding and interaction with the protein. (**B**) Modelled structure of human USP30 in complex with **USP30-I-1**. Structure of human USP30 catalytic domain highlighting the modelled position of **USP30-I-1** shown as a stick representation with carbon atoms colored green. The thumb, palm and fingers subdomains of the catalytic domain and the catalytic cysteine, Cys77, are highlighted. Regions identified in the HDX-MS analysis of USP30 in the presence of **USP30-I-1** are colored red (mean perturbation over 60 min of 5-20%) and magenta (mean perturbation over 60 min of <60%). (**C**). Close-up view of the putative **USP30-I-1** binding site highlighting flanking residues and key hydrogen-bonding interactions represented as dotted lines. The positions of blocking loop 1 (BL1), blocking loop 2 (BL2), and the switching loop (SL) are highlighted. Figure prepared using PyMOL (The PyMOL Molecular Graphics System, version 2.5.8; Schrödinger, LLC).

Finally, we performed molecular docking studies to computationally validate our *in vitro* work. As illustrated in the pose displayed in **Figures 3B** and **3C**, **USP30-I-1** is predicted to bind in the thumb-palm cleft that guides the ubiquitin C-terminus into the active site. **USP30-I-1** forms a thioimidate with the catalytic cysteine, Cys77, with the imine moiety accepting a hydrogen bond from the side chain of Asn72 and acting as a hydrogen bond donor to the main chain carbonyl of Gly74. In addition, the amide moiety of **USP30-I-1** hydrogen bonds to the main chain carbonyl of Gln160 (**Figure 3C**). The docking pose for **USP30-I-1** correlates perfectly with our HDX-MS results, with both suggesting that the USP30 catalytic Cys77 is the primary binding site of **USP30-I-1.** In addition, the docking pose of **USP30-I-1** correlates well with the X-ray structures of human USP30 catalytic domain in complex with the covalent inhibitors, **552** (PDB code: 8D1T; unpublished) and **829** (PDB code: 8D0A; unpublished) (**Figures 4A, 4B, and 4C**). In the complexes of USP30 with **552** and **829**, the switching loop adopts an “in” conformation, whereas in the complex with ubiquitin-propargylamide (PDB code: 5OHK ^36^) it adopts an “out” conformation, which, coupled with conformational differences in blocking loops 1 and 2, allow the C-terminal tail of the ubiquitin substrate to be accommodated in the thumb-palm cleft leading to active site.

**Figure 4.**
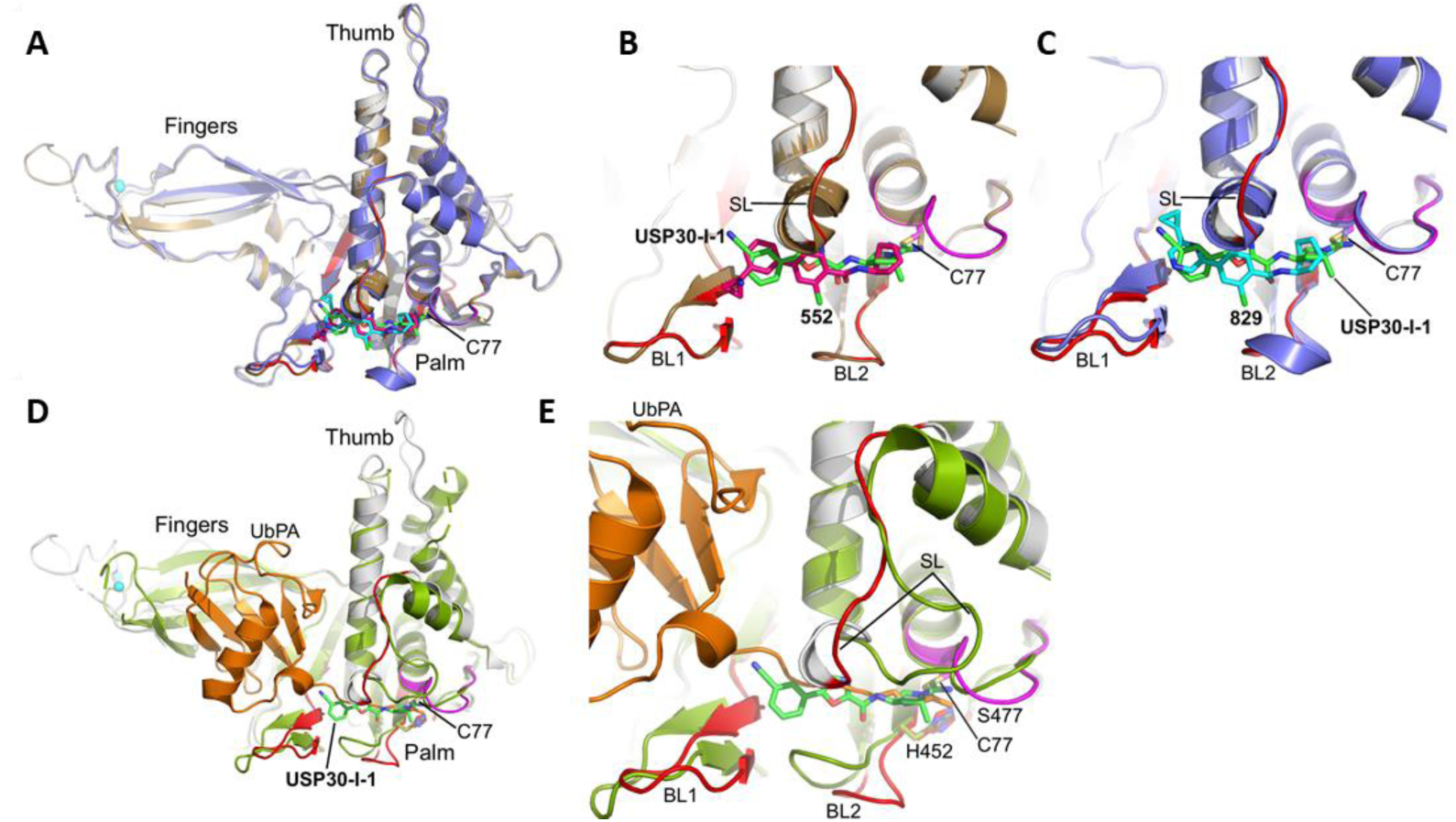
**Comparison of USP30-I-1 binding to USP30 with Ub and analogous covalent USP30 inhibitors**. (**A**). Superposition of the modelled structure of human USP30 in complex with **USP30-I-1** on the X-ray structures of human USP30 in complex with the covalent inhibitors, **552** (PDB code: 8D1T) and **829** (PDB code: 8D0A). The thumb, palm and fingers subdomains of the catalytic domain and the catalytic cysteine, Cys77, are highlighted. The catalytic domain of the modelled structure of human USP30 in complex with **USP30-I-1** is colored grey with regions implicated in compound binding from the HDX-MS analysis colored red (mean perturbation over 60 min of 5-20%) and magenta (mean perturbation over 60 min of <60%). **USP30-I-1** (carbon atoms in green), **552** (carbon atoms in hot pink), and **829** (carbon atoms in cyan) are shown as stick representations. The catalytic domain of USP30 in complex with **552** and **829** are colored brown and violet, respectively. The modelled pose of **USP30-I-1** correlates well with **552** and **829**. (**B**). Close-up view of the superimposed structure of human USP30 in complex with **552** (carbon atoms in hot pink) with the modelled structure of **USP30-I-1** (carbon atoms in green). The positions of BL1, BL2, and SL are highlighted. (**C**). Close-up view of the superimposed structure of human USP30 in complex with **829** (carbon atoms in cyan) with the modelled structure of **USP30-I-1** (carbon atoms in green). The positions of BL1, BL2, and SL are highlighted. (**D**). Superposition of the modelled structure of human USP30 in complex with **USP30-I-1** on the X-ray structure of human USP30 in complex with ubiquitin-propargylamide (UbPA; PDB code: 5OHK). The catalytic domain of the modelled structure of human USP30 in complex with **USP30-I-1** is colored grey with regions implicated in compound binding from HDX-MS analysis are colored as above. **USP30-I-1** is shown with carbon atoms colored green. The catalytic domain of USP30 in complex with UbPA is colored lime with UbPA shown in orange. **USP30-I-1** is predicted to bind in the thumb-palm cleft that guides the ubiquitin C-terminus into the active site. (**E**). Close-up view of the superimposed structures of human USP30 in complex with UbPA and the modelled structure of **USP30-I-1** (carbon atoms in green). Catalytic triad residues (C77, H452 and S477) are highlighted and shown as stick representations. Figure prepared using PyMOL (The PyMOL Molecular Graphics System, version 2.5.8; Schrödinger, LLC).

Hence, the predicted binding site of **USP30-I-1** based on our modelling studies would sterically clash with the C-terminal tail of ubiquitin (**Figures 4D** and **4E**).

Distinct apo-form and ubiquitin-bound conformational states have been observed for other USP family members and are implicated in the catalytic cycle ^37, 38^. In addition, several USP inhibitors have been shown to preferentially target the apo-form conformation. For example, the non-covalent USP7 inhibitor, **FT671** (PDB code: 5NGE) and covalent USP7 inhibitor, **FT827** (PDB code: 5NGF), specifically target the catalytically incompetent apo-form state in which the switching loop adopts an “in” conformation ^39^. A comparison of **USP30-I-1** with **FT827** reveals that the binding site is broadly similar but that there are conformational differences in the region accommodating the catalytic cysteine, the switching loop, and blocking loops 1 and 2 (**Figures 5A and 5B**). These conformational differences result in **FT827** extending closer towards the fingers subdomain than the modelled position of **USP30-I-1**. We envisage that USP30 may also adopt distinct conformational states during its catalytic cycle, which can be exploited by inhibitors, but await the determination of an experimental structure of apo-form USP30 catalytic domain to confirm our hypothesis.

**Figure 5.**
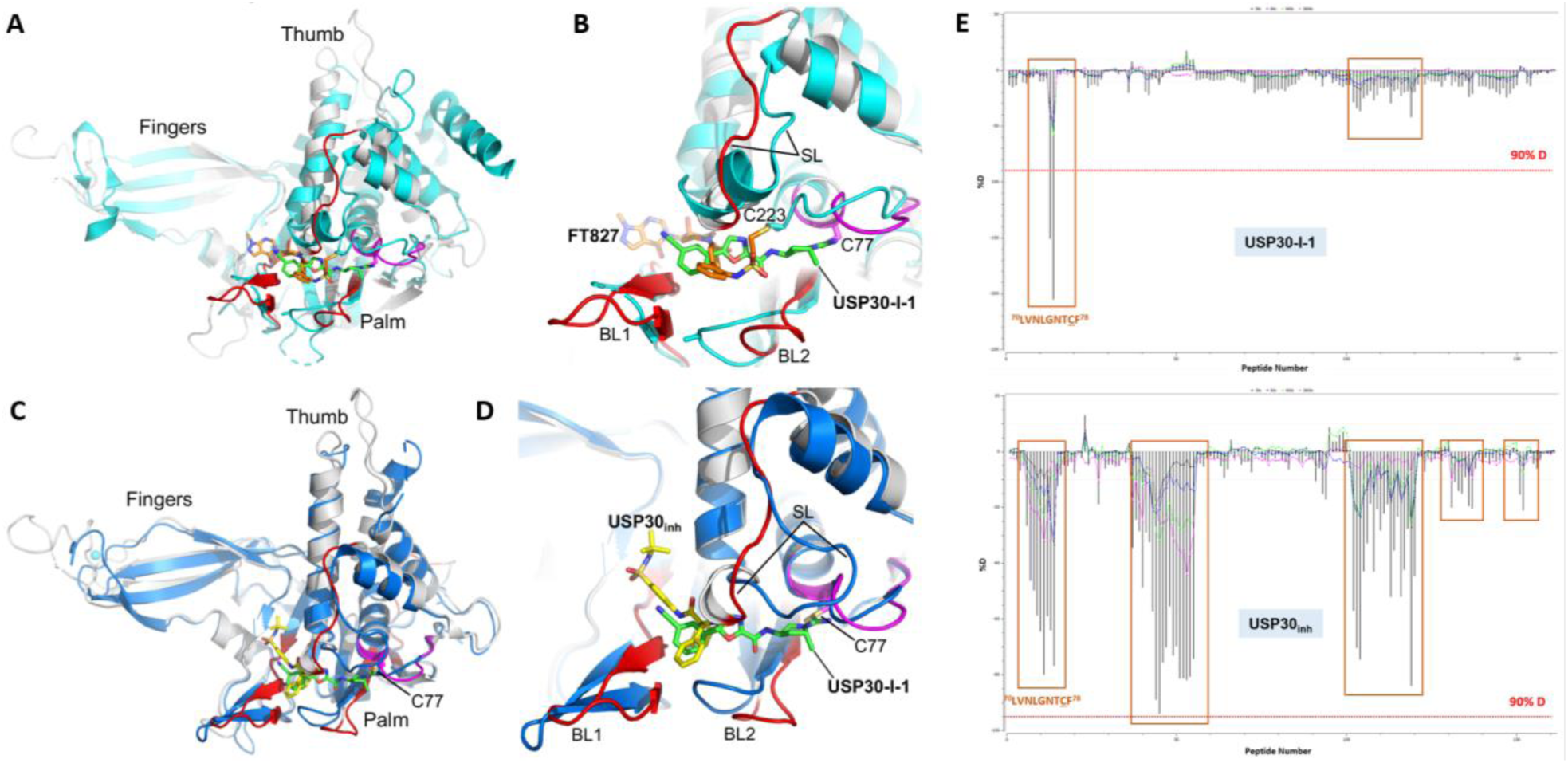
Comparison of covalent and non-covalent inhibitor binding in USP7 and USP30. (**A**) Superposition of the modelled structure of human USP30 in complex with **USP30-I-1** on the X-ray structure of human USP7 in complex with the covalent inhibitor, **FT827** (PDB code: 5NGF; r.m.s.d.= 2.5 Å, 249 residues aligned). (**B**) Close-up view of the superimposed modelled structure of human USP30 in complex with **USP30-I-1** (green carbon atoms) on the X-ray structure of human USP7 in complex with the covalent inhibitor, **FT827** (orange carbon atoms). The positions of BL1, BL2, SL, and the catalytic cysteines (Cys77 in USP30 and Cys223 in USP7) are highlighted. The Cαs of Cys77 and Cys223 are separated by approximately 5.2 Å, which, combined with differences in the conformations of BL1, BL2 and SL, result in **FT827** extending closer towards the fingers subdomain than **USP30-I-1**. (**C**) Superposition of the modelled structures of human USP30 in complex with **USP30-I-1** and **USP30**_inh_ ^24^. The catalytic domain of the modelled structure of human USP30 in complex with **USP30-I-1** is colored grey with regions implicated in compound binding from HDX-MS analysis colored red (mean perturbation over 60 min of 5-20%) and magenta (mean perturbation over 60 min of <60%). **USP30-I-1** is shown with carbon atoms colored green. The catalytic domain of the modelled structure of human USP30 in complex with **USP30**_inh_ is colored blue. **USP30**_inh_ is shown with carbon atoms colored yellow. (**D**) Close-up view of the superimposed modelled structures of human USP30 in complex with **USP30-I-1** (carbon atoms in green) and **USP30**_inh_ (yellow carbon atoms). The positions of BL1, BL2, and SL are highlighted. Both inhibitors are predicted to bind within the thumb-palm cleft with **USP30**_inh_ residing approximately 7.9 Å away from Cys77 and extending out towards the fingers subdomain. (**E**) HDX-MS residuals plot of USP30 in complex with **USP30-I-1** and **USP30**_inh_. A greater overall solvent protection is observed for USP30 in the presence of the non-covalent inhibitor, as compared to its covalent counterpart. **USP30-I-1** primarily induces solvent protection in the region encompassing the catalytic Cys77, whereas the non-covalent **USP30**_inh_ results in smaller HDX-MS perturbations, albeit, extended to several regions of the protein. Figure prepared using PyMOL (The PyMOL Molecular Graphics System, version 2.5.8; Schrödinger, LLC).

There are 26 residues with an atom residing within 5Å of the predicted **USP30-I-1** binding site (**Figure 3B**). Inhibitor selectivity is likely to be conferred by sequence substitutions in these residues compared with other USP family members, coupled with conformational differences in the regions flanking the proposed **USP30-I-1** binding site including the switching and blocking loops. In summary, our molecular docking analysis identified residues implicated in **USP30-I-1** binding, which is in perfect agreement with those identified through differential HDX-MS.

### Comparison of USP30 inhibition by covalent cyanopyrrolidine USP30-I-1 and non-covalent benzosulfonamide USP30_inh_ compound

We previously performed biochemical, kinetic, and structural characterization of USP30 in complex with a non-covalent small-molecule benzosulfonamide **USP30**_inh_ ^24^. By directly comparing our two studies, we could decipher commonalities and differences between covalent and non-covalent mechanisms of USP30 inhibition in terms of the kinetics of complex formation and structural dynamics.

For covalent compounds, IC_50_ is a poor measure of potency as it is a time-dependent process and often correlates poorly with efficacy. Determination of k_inact_/K_I_ is the gold standard for assessing the potency of covalent inhibitors. k_inact_/K_I_ is a second order rate constant which defines the efficiency of covalent bond formation by incorporating the affinity of the initial reversible binding of the inhibitor (K_I_) and the rate at which the enzyme is inactivated by covalent bond formation (k_inact_) with a higher value corresponding to a more potent inhibitor. **USP30-I-1** shows both a high initial binding affinity (K_I_ ∼ 350 nM) and a fast rate of specific inactivation (k_inact_ ∼ 0.15 s^-1^). Both **USP30-I-1** and non-covalent **USP30**_inh_ show time-dependent inhibition, with **USP30**_inh_ showing two-step slow, tight binding kinetic behavior consistent with a covalent inhibitor. The second step of inhibition by **USP30**_inh_ is essentially irreversible (k_6_ = 0.00033s^-1^) allowing us to compare k_5_/K_iapp_ for **USP30**_inh_ with k_inact_/K_I_ for **USP30-I-1** as a measure of potency. Using these values, the compounds have similar potency (0.18 μΜ^-1^s^-1^ and 0.20 μM^-1^s^-1^ for **USP30-I-1** and **USP30**_inh_ respectively), in the case of **USP30**_inh_ the potency is driven by the rate at which the irreversible complex forms with initial affinity being lower than that of **USP30-I-1** (1.27 μM and 350 nM for **USP30-I-1** and **USP30**_inh_ respectively). It will be interesting to see whether these differences have an effect on cellular potency.

Our previous study also employed HDX-MS and computational docking to elucidate the molecular architecture and geometry of USP30 complex formation with the non-covalent inhibitor, **USP30**_inh_ ^24^. Both compounds interact with the catalytic Cys77, but **USP30-I-1** induces a higher degree of solvent protection than **USP30**_inh_. Only the catalytic region undergoes substantial perturbation in the presence of the covalent compound, suggesting that it is a more targeted interaction. A much larger area is perturbed in the presence of the non-covalent inhibitor, implying that the compound is moving about and reorienting itself within the binding pocket (**Figures S4** and **5E**). **USP30**_inh_ is also predicted to bind to the thumb-palm cleft of the catalytic domain. However, compared with **USP30-I-1**, which forms a covalent adduct with the catalytic Cys77, the modelled position of **USP30**_inh_ resides approximately 7.9 Å away at its closest point from the thiol side chain of this cysteine (**Figures 5C** and **5D**). The X-ray structure of human USP30 catalytic domain in complex with UbPA (PDB code: 5OHK; Gersch *et al*., 2017) was used as the target receptor for **USP30**_inh_ since there were no inhibitor-bound structures available at that time, which results in differences in conformation in the switching and blocking loop regions compared with the structure of USP30 in complex with the covalent inhibitor, **552** (PDB code: 8D1T), that was used for **USP30-I-1** docking studies. However, we note that the docking pose of **USP30**_inh_ correlated well with the HDX-MS data and is predicted to bind to an equivalent site as the non-covalent USP7 inhibitor, **FT671**, which resides approximately 5 Å away from the catalytic cysteine (Turnbull *et al*., 2017). Hence, whilst both studies indicate that **USP30**_inh_ and **USP30-I-1** are likely to reside within the thumb-palm cleft, differences in their HDX-MS profiles suggest that there are differences in their binding modes, which places **USP30**_inh_ closer towards the fingers subdomain compared with the predicted binding site for **USP30-I-1** (**Figures 5E**).

## CONCLUSIONS

**USP30-I-1** represents a covalent, small-molecule cyanopyrrolidine inhibitor, which exhibits high potency and selectivity for the mitochondrial DUB, USP30. HDXMS studies reveal that covalent attachment of **USP30-I-1** to Cys77 results in significant solvent protection in the region that flanks the catalytic cysteine whereas less noticeable structural and conformational changes are seen in other regions. Inhibitor binding blocks the interaction of USP30 with Ub substrate molecules, preventing isopeptide bond cleavage. This enhanced understanding of the molecular mechanisms surrounding USP30 inhibition will aid in the creation of next-generation inhibitors targeting neurodegenerative and cardiovascular diseases.

## DATA AVAILABILITY

All ABPP-MS proteomics raw files have been deposited to the ProteomeXchange Consortium and can be accessed through the identifier PXD054041. Similarly, HDX-MS data can be downloaded using PXD054057.

## SUPPORTING INFORMATION

### Enzyme Kinetics

The traditional method to determine the kinetics of a covalent compound was also used. Progress curves were fitted to Eq.2, where [P] is product formed, *t* is time, *v*_i_ is the initial rate of the reaction and k_obs_ the rate at which the system is inactivated.

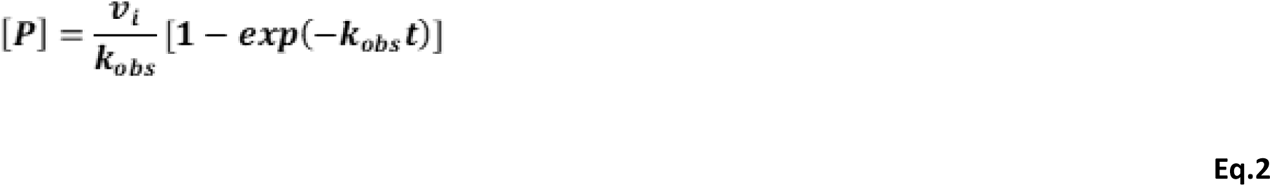

The resulting k_obs_ values were plotted against concentration and fitted to Eq.3a to obtain K_I_ and k_inact_.

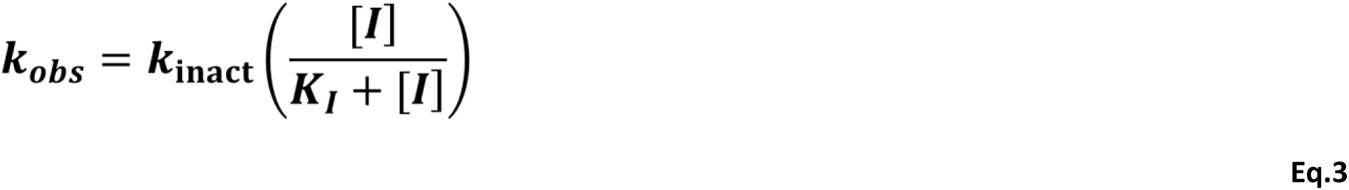

**Figure S1. Purity profile of USP30-I-1.** The compound was deemed 100% pure by HPLC analysis and when measured at 254 nm.

**Figure S2. ABPP-MS analysis of USP30-I-1 for inhibiting USP30.** LC-MS/MS quantitation of HA-Ub-PA enriched DUBs with **USP30-I-1** at 0 and 10 Μm relative to positive control (****p < 0.0001).

**Figure S3. Comparison of ABPP-MS of USP30-I-1 for inhibiting USP30 and USP10. A.** No strong HA-Ub-PA labeling of USP10. **B.** LC-MS/MS quantitation of HA-Ub-PA enriched USP30 & USP10, demonstrating higher potency of **USP30-I-1** for USP30 inhibition compared with USP10.

**Figure S4. HDX-MS Volcano Plots of USP30 in complex with USP30-I-1 and USP30**_inh_. A greater overall solvent protection is observed for USP30 in the presence of the non-covalent inhibitor, as compared to its covalent counterpart. **USP30-I-1** primarily induces solvent protection in the region encompassing the catalytic Cys77.

## AUTHOR INFORMATION

### Corresponding Authors

**Darragh P O’Brien** – Structural and Mechanistic Proteomics Laboratory, Target Discovery Institute, Centre for Medicines Discovery, Nuffield Department of Medicine, University of Oxford, Oxford OX3 7FZ, United Kingdom; ORCID: 0000-0003-4924-7795;

**Andrew P Turnbull** – Cancer Research Horizons, Francis Crick Institute, 1 Midland Road, London NW1 1AT, United Kingdom;

**Benedikt M Kessler** – Biological Mass Spectrometry Laboratory, Target Discovery Institute, Centre for Medicines Discovery, Nuffield Department of Medicine, University of Oxford, Oxford OX3 7FZ, United Kingdom; ORCID: 0000-0002-8160-2446;

### Authors

**Hannah BL Jones** – Target Discovery Institute, Centre for Medicines Discovery, Nuffield Department of Medicine, University of Oxford, Oxford OX3 7FZ, United Kingdom

**Yuqi Shi** – Thermo Fisher Scientific, San Jose, California, USA

**Franziska Guenther** – ARUK-Oxford Drug Discovery Institute, Centre for Medicines Discovery, Nuffield Department of Medicine, University of Oxford, Oxford OX3 7FZ, United Kingdom

**Iolanda Vendrell** – Target Discovery Institute, Centre for Medicines Discovery, Nuffield Department of Medicine, University of Oxford, Oxford OX3 7FZ, United Kingdom

**Rosa Viner** – Thermo Fisher Scientific, San Jose, California, USA

**Paul E Brennan** – ARUK-Oxford Drug Discovery Institute, Centre for Medicines Discovery, Nuffield Department of Medicine, University of Oxford, Oxford OX3 7FZ, United Kingdom

**Emma Mead** – ARUK-Oxford Drug Discovery Institute, Centre for Medicines Discovery, Nuffield Department of Medicine, University of Oxford, Oxford OX3 7FZ, United Kingdom

**Tryfon Zarganes-Tzitzikas** – ARUK-Oxford Drug Discovery Institute, Centre for Medicines Discovery, Nuffield Department of Medicine, University of Oxford, Oxford OX3 7FZ, United Kingdom

**John B Davis** – ARUK-Oxford Drug Discovery Institute, Centre for Medicines Discovery, Nuffield Department of Medicine, University of Oxford, Oxford OX3 7FZ, United Kingdom

**Adán Pinto-Fernández** – Target Discovery Institute, Centre for Medicines Discovery, Nuffield Department of Medicine, University of Oxford, Oxford OX3 7FZ, United Kingdom; Chinese Academy of Medical Sciences Oxford Institute, University of Oxford, UK

**Katherine S England** – ARUK-Oxford Drug Discovery Institute, Centre for Medicines Discovery, Nuffield Department of Medicine, University of Oxford, Oxford OX3 7FZ, United Kingdom

**Emma J Murphy** – ARUK-Oxford Drug Discovery Institute, Centre for Medicines Discovery, Nuffield Department of Medicine, University of Oxford, Oxford OX3 7FZ, United Kingdom

### Author contributions

H. B. L. J. and A.P.F. performed the ABPP-MS experiments and data analysis. F.G. and E.J.M. performed enzyme kinetics and data analysis. D. P. O., Y.S., and R.V. performed the HDX-MS experiments and data analysis. I. V. performed the ABPP-MS data acquisition. K.S.E. and A.P.T. performed the computational docking. P. E. B., E. M., T.Z.T., J. B. D., A. P. T., B.M.K. and A. P-F, took part in project design and discussion. D. P. O., H. B. L. J., Y.S., K.S.E., E.J.M., and A.P.T. prepared the first draft of the manuscript. All authors edited and approved the final manuscript.

### Notes

The authors declare that they have no conflicts of interest, financial or otherwise, with the contents of this article.

## ACKNOWLEDGEMENTS

**USP30-I-1** was a kind gift from Bristol Myers Squibb. We greatly appreciate the input of Daryl S Walter from Evotec and Jeff Schkeryantz from Bristol Myers Squibb over the duration of the project. Our tremendous gratitude goes to Alzheimer’s Research UK for their sustained, long-term funding of the ARUK-Oxford Drug Discovery Institute (ODDI; grant no. ARUK-2021DDI-OX). Heartfelt thanks go to the Chinese Academy of Medical Sciences (CAMS) Innovation Fund for Medical Science (CIFMS), China (grant number: 2018-I2M-2-002) (BMK, APF), the G & K Boyes Charitable Trust, and to the late Mr & Mrs James Hardwick for their generous support of the ODDI Medicinal Chemistry Laboratory.

## ABBREVIATIONS

ABPP-MS: Activity-Based Protein Profiling Mass Spectrometry
DIA: Data Independent Acquisition
DMSO: dimethyl sulfoxide
DUB: Deubiquitinase
HDX-MS: Hydrogen Deuterium eXchange Mass Spectrometry
LFQ: Label-Free Quantitation
MOM: Mitochondrial Outer Membrane
PD: Parkinson’s Disease
PDB: Protein Data Bank
SAR: Structure Activity Relationship
Ub: Ubiquitin
UPS: Ubiquitin Proteasome System
USP30: Ubiquitin Specific Protease 30
USP30-I-1: covalent inhibitor of USP30

