## Supplementary Figures for "Structural Dynamics of the Ubiquitin Specific Protease USP30 in Complex with a Cyanopyrrolidine-Containing Covalent Inhibitor"

**FractionLynx Report -** Page 2

Sample: 2 Vial:5,3:5,H ID:
File: Date:20-Oct-2023 Time:16:24:32
Description:

Printed: Wed Nov 01 16:01:49 2023

**Sample Report (continued):**

UV Detector: 254 Smooth (SG, 1x1) 2.499e-1
Range: 3.11e-1

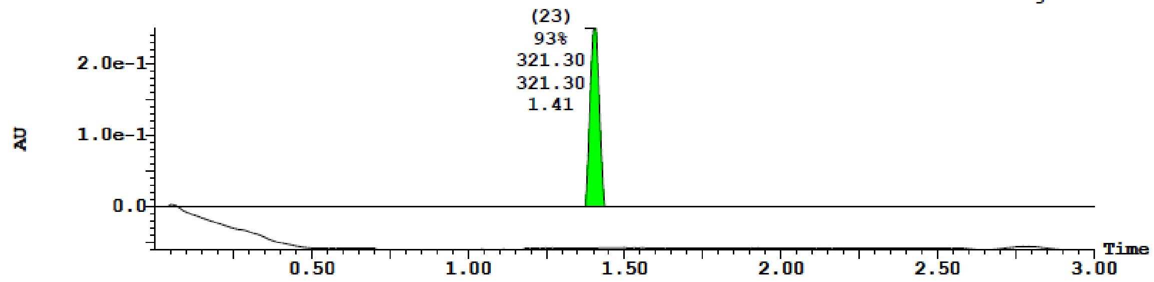


| Peak Number | Compound | Time | AreaAbs | Area %Total | Width | Height | Mass Found |
|---|---|---|---|---|---|---|---|
| 23 | Found | 1.41 | 1e+004 | 93.41 | 0 | 3e+005 | 321.30, 321.30 |
| 26 | | 1.71 | 9e+002 | 6.59 | 0 | 2e+004 | |

(2) ELSD Signal Smooth (SG, 1x1) 670.894
Range: 671.639

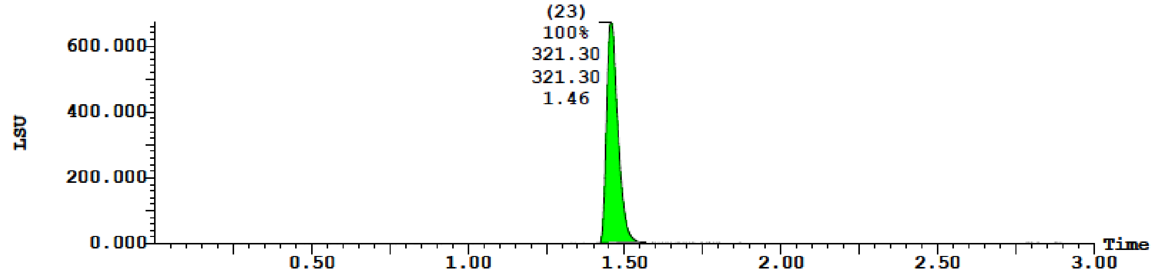


| Peak Number | Compound | Time | AreaAbs | Area %Total | Width | Height | Mass Found |
|---|---|---|---|---|---|---|---|
| 23 | Found | 1.46 | 3e+004 | 100.00 | 0 | 7e+005 | 321.30, 321.30 |

**Figure S1. Purity profile of USP30-I-1.** The compound was deemed 100% pure by HPLC analysis and measurement at 254 nm.

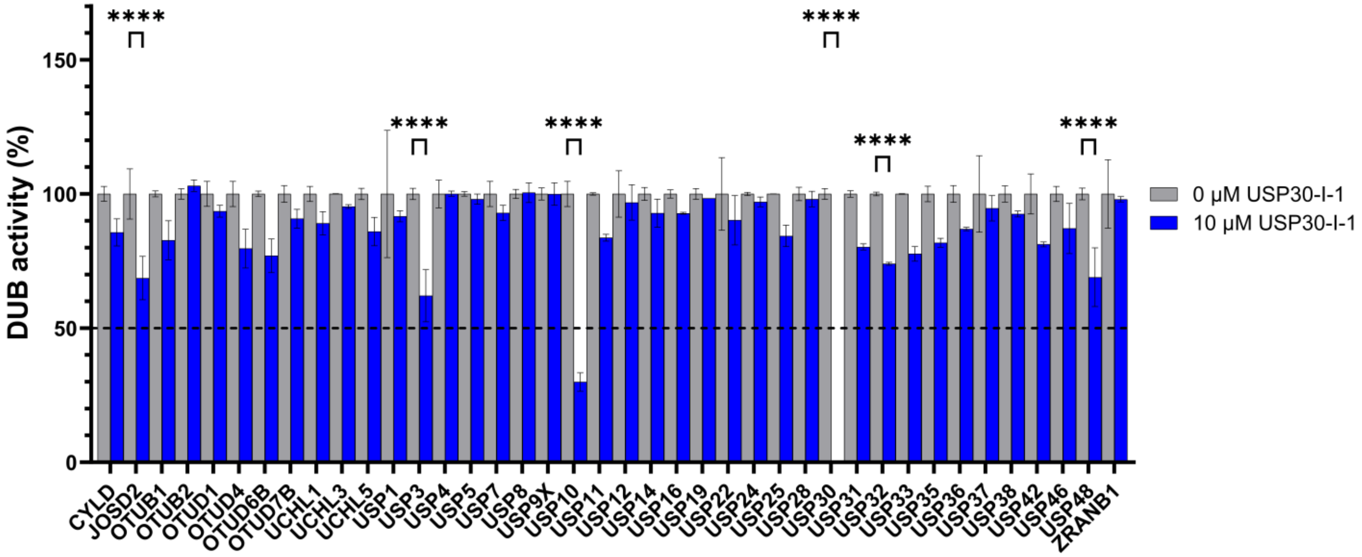


**Figure S2. ABPP-MS analysis of USP30-I-1 for inhibiting USP30. A.** LC-MS/MS quantitation of HA-Ub-PA enriched DUBs with **USP30-I-1** at 0 and 10 µM relative to positive control (****p < 0.0001).

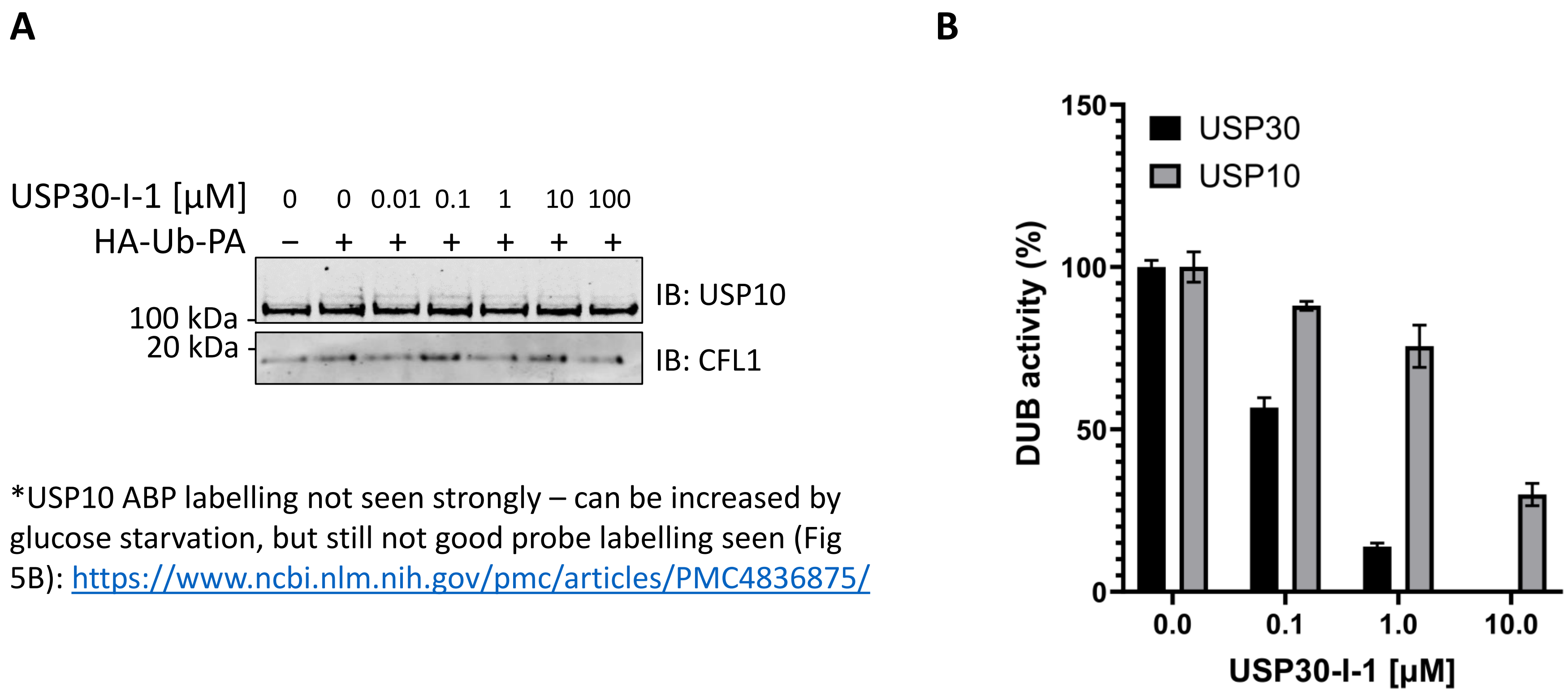


*USP10 ABP labelling not seen strongly – can be increased by glucose starvation, but still not good probe labelling seen (Fig 5B): https://www.ncbi.nlm.nih.gov/pmc/articles/PMC4836875/

**Figure S3. Comparison of ABPP-MS of USP30-I-1 for inhibiting USP30 and USP10. A.** No strong HA-Ub-PA labelling of USP10. **B.** LC-MS/MS quantitation of HA-Ub-PA enriched USP30 & USP10, demonstrating higher potency of USP30-I-1 for USP30 inhibition compared with USP10.

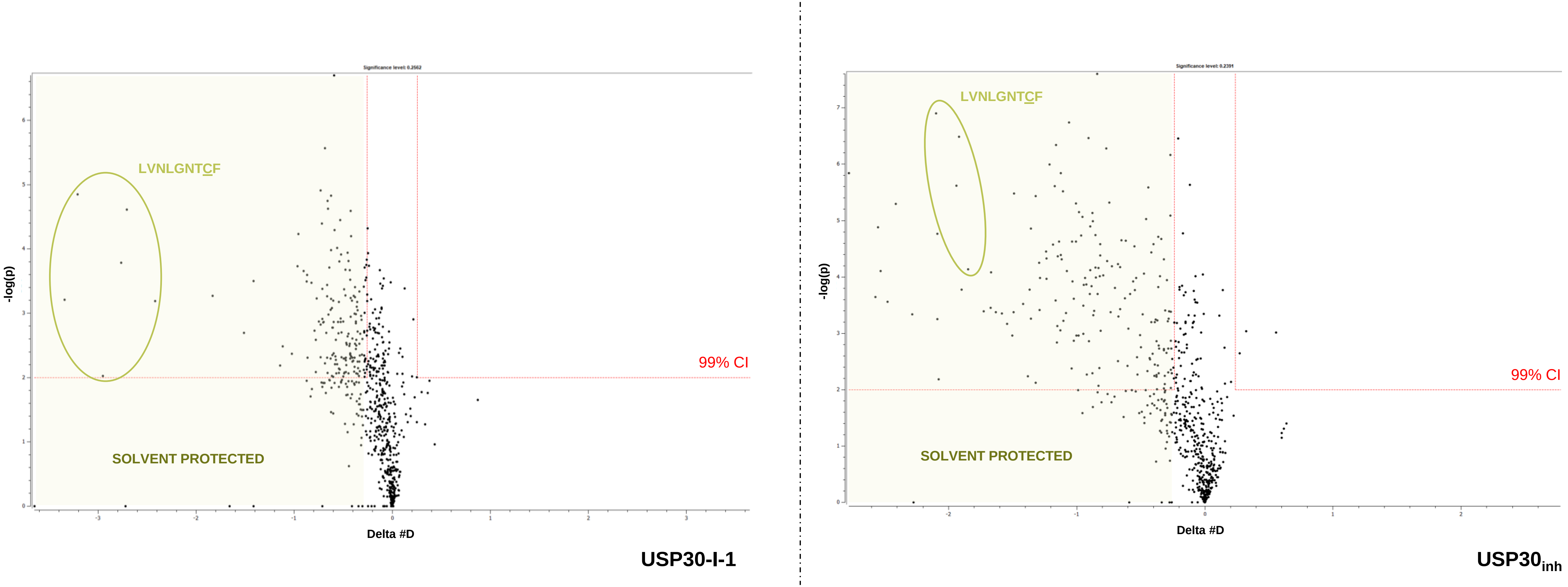


**Figure S4. HDX-MS volcano plots of USP30 in complex with USP30-I-1 and USP30$_{inh}$.** A greater overall solvent protection is observed for USP30 in the presence of the non-covalent inhibitor, as compared to its covalent counterpart. **USP30-I-1** primarily induces solvent protection in the region encompassing the catalytic Cys77.
